# Making many out of few: deep generative models for single-cell RNA-sequencing data

**DOI:** 10.1101/2020.05.27.119594

**Authors:** Martin Treppner, Adrián Salas-Bastos, Moritz Hess, Stefan Lenz, Tanja Vogel, Harald Binder

**Affiliations:** Institute of Medical Biometry and Statistics, Faculty of Medicine and Medical Center - University of Freiburg, Freiburg, 79104, Germany; Freiburg Center of Data Analysis and Modelling, Mathematical Institute - Faculty of Mathematics and Physics, University of Freiburg, Freiburg, 79104, Germany; Institute of Anatomy and Cell Biology, Department of Molecular Embryology, Medical Faculty, University Freiburg, Freiburg, 79104, Germany; Faculty of Biology, University of Freiburg, Freiburg, Germany; Center for Basics in NeuroModulation (NeuroModul Basics), University of Freiburg, Freiburg, 79104, Germany; Freiburg Institute for Advanced Studies (FRIAS), University of Freiburg, Germany

## Abstract

Deep generative models, such as variational autoencoders (VAEs) or deep Boltzmann machines (DBM), can generate an arbitrary number of synthetic observations after being trained on an initial set of samples. This has mainly been investigated for imaging data but could also be useful for single-cell transcriptomics (scRNA-seq). A small pilot study could be used for planning a full-scale study by investigating planned analysis strategies on synthetic data with different sample sizes. It is unclear whether synthetic observations generated based on a small scRNA-seq dataset reflect the properties relevant for subsequent data analysis steps.

We specifically investigated two deep generative modeling approaches, VAEs and DBMs. First, we considered single-cell variational inference (scVI) in two variants, generating samples from the posterior distribution, the standard approach, or the prior distribution. Second, we propose single-cell deep Boltzmann machines (scDBM). When considering the similarity of clustering results on synthetic data to ground-truth clustering, we find that the *scVI*_*posterior*_ variant resulted in high variability, most likely due to amplifying artifacts of small data sets. All approaches showed mixed results for cell types with different abundance by overrepresenting highly abundant cell types and missing less abundant cell types. With increasing pilot dataset sizes, the proportions of the cells in each cluster became more similar to that of ground-truth data. We also showed that all approaches learn the univariate distribution of most genes, but problems occurred with bimodality. Overall, the results showed that generative deep learning approaches might be valuable for supporting the design of scRNA-seq experiments.

## Introduction

Deep generative models, such as variational autoencoders (VAEs)^1,2^ or deep Boltzmann machines (DBMs)^3^, can learn the joint distribution of various types of data, and impressive results have been obtained, e.g., for generating super-resolution images in microscopy^4^ and more generally for imputation^5,6^. This raises the question of whether such techniques could also be trained on data with a rather small number of samples, e.g., obtained from pilot experiments, for subsequently generating larger synthetic datasets. Such synthetic observations could, in particular, inform the design of single-cell RNA sequencing (scRNA-seq) experiments, e.g., by exploring planned subsequent analysis steps, such as clustering, on synthetic datasets of different sizes.

ScRNA-seq experiments result in data reflecting gene expression for individual cells in tissues, leading to an improved understanding of cell-type composition. Due to the underlying complexity of the data, deep generative approaches are increasingly used to investigate the structure of scRNA-seq data by learning a low-dimensional latent representation of gene expression within cells. Often, the focus of these applications is on exploring latent features of the data —representing cell types— after which they are used for clustering, imputation, or differential expression analysis^6–8^. As indicated, another interesting property of these generative approaches is that they can provide synthetic data once trained on some dataset. However, the quality of the data generated from these models is challenging to evaluate and requires cautious examination, depending on the field of application and the research question^9^. In the following, we investigate the quality of synthetic data from generative models and illustrate various obstacles for its application based on an example of the design of single-cell RNA-seq experiments.

As the experimental design of scRNA-seq studies is often based on simulations^10–14^, synthetic data could be useful, e.g., when training a generative approach on some pilot data. Sampling from latent representations of generative models then allows for generating in-silico expression patterns, ideally reflecting the most important patterns from the pilot data, and can subsequently be utilized for planning experiments. More specifically, researchers would specify different numbers of cells to be simulated, then apply downstream analyses to the simulated data, after which they evaluate the number of cells needed for detecting patterns of interest, such as clusters comprising rare cell types.

To investigate the authenticity of the synthetic data using experimental design as an example, we follow the procedure shown in **Figure 1**. First, we extract small pilot datasets from the larger original data by subsampling (**Figure 1 A**). Next, we train the deep generative models on the subsampled pilot datasets and generate synthetic data in the size of the original study (**Figure 1 B**). We apply downstream analyses to both the original data and the synthetic data (**Figure 1 C**), after which we examine the quality of the synthetic observations using various evaluation approaches (**Figure 1 D**).

**Figure 1.**
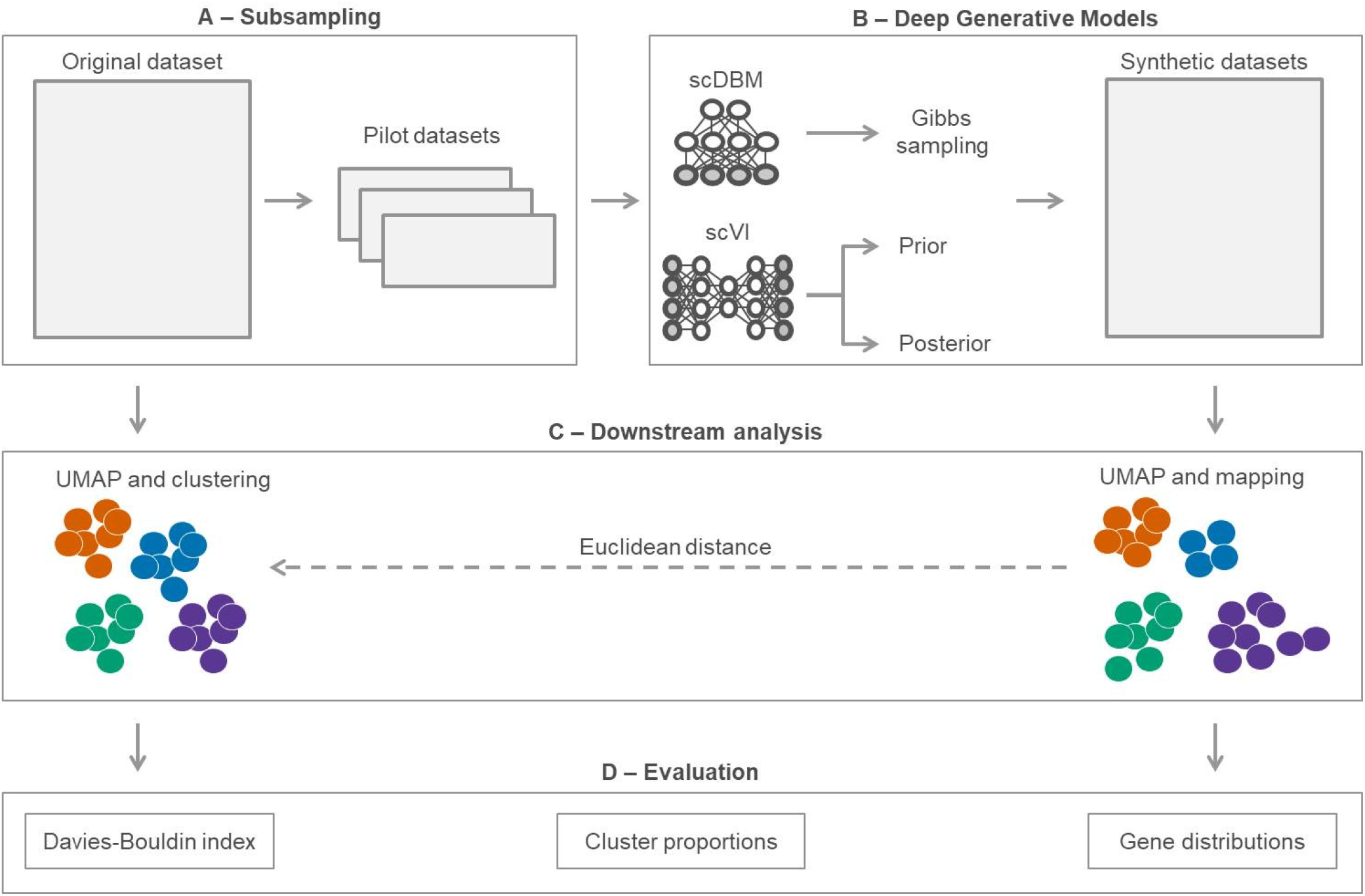
Design for evaluating the performance of deep generative models with small pilot datasets: (A) Take a subsample from an original dataset to obtain pilot data with known ground truth. (B) Train the deep generative approaches on the pilot dataset and generate synthetic data in the original data size. (C) Apply dimensionality reduction with UMAP and Seurat clustering to the original data and map each synthetic observation to the closest observation from the original data, thus getting a cluster assignment. (D) Evaluate the quality of synthetic samples based on the Davies-Bouldin index, cluster proportions, and distributions per gene. The complete analysis is performed for different sizes of pilot datasets (384, 768, 1152, 1536, 1920, and 2304 cells) and repeated 30 times for each size.

While VAEs have already been proposed for scRNA-seq data^7^, DBMs still need to be adapted. We show how this can be achieved using a negative binomial distribution and incorporating a regularized Fisher scoring algorithm to estimate the inverse dispersion parameter. We chose DBMs because synthetic observations are generated by Gibbs sampling, which has theoretical properties that are potentially advantageous for working with smaller sample sizes than variational inference in VAEs^15,16^.

VAEs reconstruct their input through a bottleneck layer that corresponds to a low-dimensional latent representation. They offer two ways of generating samples from the latent representation. Most commonly, samples are generated from the posterior, which is the latent variables’ probability given the original data. In a pilot study setting, this will typically mean that multiple copies of the original observations have to be used to obtain a larger synthetic dataset. This might lead to an amplification of sampling bias, as patterns or random fluctuations from single cells could be over-emphasized. In contrast, sampling from the prior might produce samples from a diverse region of the latent space. In our evaluation together with DBMs, we therefore not only investigate the performance of VAEs when feeding in the original data multiple times for obtaining a larger number of cells but also when sampling directly from the prior, which has —to our knowledge— not been considered in the scRNA-seq literature so far.

## Results

To examine the quality of the data generated by single-cell variational inference (scVI) and single-cell DBMs (scDBMs), we used the example of designing a scRNA-seq experiment. By mimicking a situation where we want to plan an experiment using a pilot study with a small number of cells, we investigated the impact of varying amounts of cells and generative approaches on the clustering performance, measured by the Davies-Bouldin index. We took 30 subsamples of 384, 768, 1152, 1536, 1920, and 2304 cells of the original dataset, trained the scDBM and scVI on these subsamples, and generated synthetic data. More precisely, we sampled from the scDBM using Gibbs sampling and from scVI using the prior and posterior distribution, respectively. The subsamples’ size is based on the number of cells that can be captured by a 384-well plate, which allows us to get an indication of the required number of plates for a sequencing experiment. We then applied UMAP and acquired the cluster labels by mapping the synthetic observations to the original data based on the Euclidean distance (**Figure 1 C**).

We have also added a performance baseline, which mainly provides an upper bound for the sampling bias. We generated negative binomial noise by randomly drawing the scale and shape parameters of a gamma distribution from Uniform(0,5) and Uniform(0,10), respectively. We used the values drawn for each cell as rate parameters in the Poisson distribution and added the resulting values to the subsampled pilot data. To be more precise, we have merged several noisy pilot datasets to achieve the original dataset’s size. For example, for a pilot dataset with 384 cells and an original dataset of 3840 cells, we replicated the noisy dataset ten times and then merged them. One would expect the Davies-Bouldin Index (DBI, see Methods) to be very high in a scenario with small pilot datasets since potential artifacts could be strongly amplified.

The results show that *scVI*_*posterior*_ exhibits high variability, especially with small datasets. In contrast, the variability for scDBM and *scVI*_*prior*_ is much lower. Regarding the variability, and thus the dependence of the models on the representativeness of the pilot data, we found similar results in other datasets (Supplementary **Figure 1, 2**). With its high variability and some extreme outliers, *scVI*_*posterior*_ sometimes even leads to worse results than our simple baseline, which is based solely on noisy pilot datasets (**Figure 2**). These findings support our hypothesis that posterior sampling in scVI leads to an amplification of the sampling bias when drawing conclusions for larger datasets based on small pilot studies.

**Figure 2.**
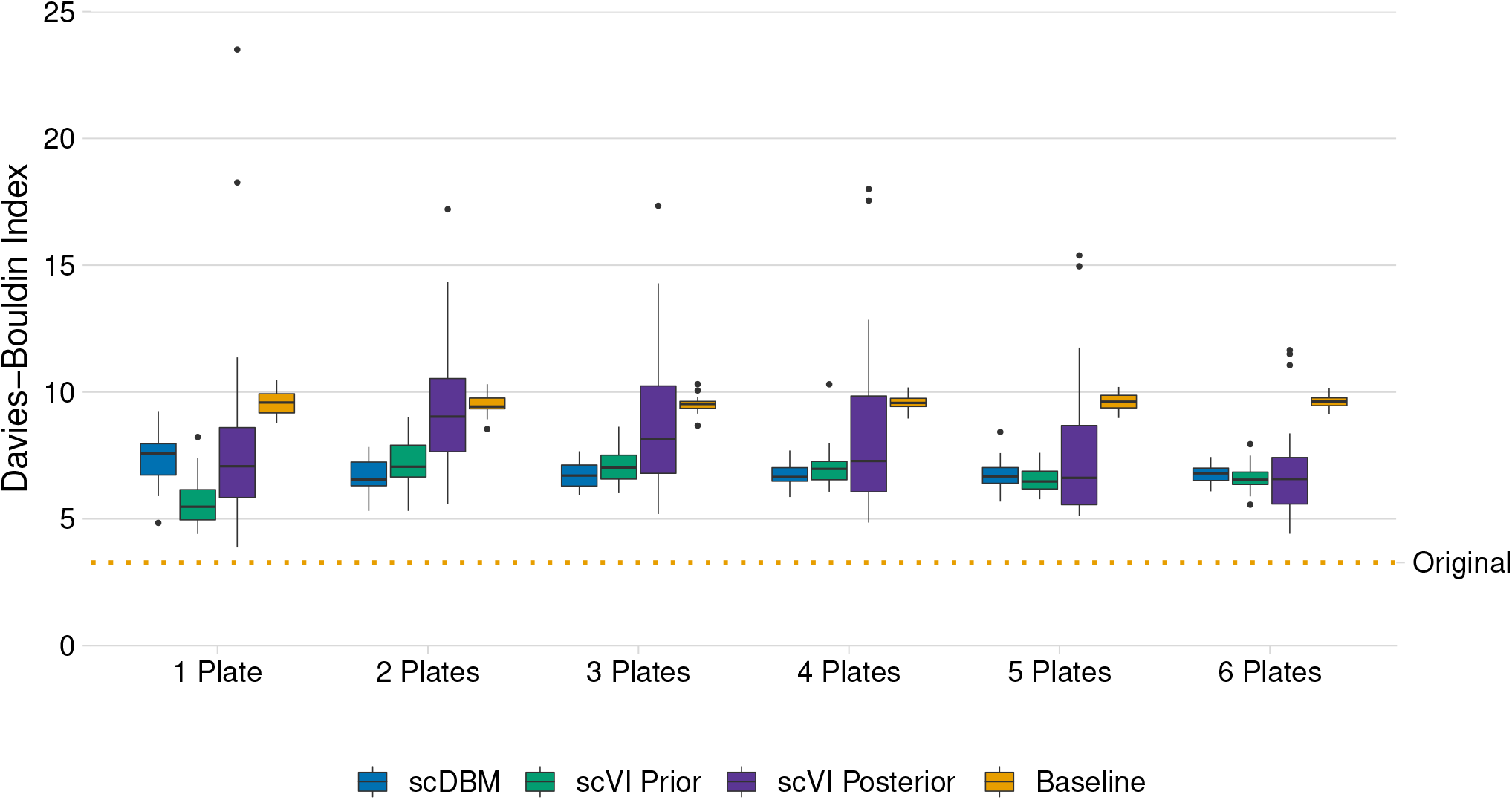
Davies-Bouldin index (DBI), indicating the quality of synthetic data generated by scDBM, scVI (prior and posterior sampling), and a baseline from pilot data of different sizes (*PBMC*4*k*). Each boxplot represents 30 subsamples from the original data (lower and upper hinges correspond to the 25th and 75th percentiles). The orange line indicates the reference DBI for the Seurat clustering on the original data with 4182 cells.

We noticed that the DBI was lowest for the smallest sample size, after which the DBI for two plates showed an increase for scVI. A possible explanation for this behavior would be that with only 384 cells drawn, some clusters are rarely or not at all represented by the sample. Since the deep generative models have a tendency to sparsity in the lower-dimensional latent space, they learn the structure with fewer clusters than they have in the original dataset. When doubling the sample size from one plate to two plates, some cells seem to be found for each cluster, but the number of cells for the models does not seem sufficient to properly learn the structure. This leads to an increase in the DBI. If the number of cases is further increased, the number of cells per cluster will also increase, and models will discover these clusters more easily.

Next, we inspected whether the models accurately estimated the proportions of cells per cluster to uncover heterogeneity and subpopulation frequencies. We have evaluated the proportions, both for each cluster and for each size of the pilot dataset. The arrows in the figure show that as the pilot data set grows larger, the ratios tend to be closer to 1 in most cases (**Figure 3**). This indicates that larger datasets might be needed to properly generate synthetic observations with adequate cluster proportions. We also inspected the marginal distributions of several exemplary genes in samples from scVI and scDBM and compared them with the distributions in the original data (**Figure 4**). We observed that the synthetic data generated from the scDBM trained on one plate match the true distribution of many genes rather well but tends to underestimate expression counts. In contrast, *scVI*_*prior*_ and *scVI*_*posterior*_ tend to overestimate expression in many genes. All methods frequently exhibit difficulties with bimodality, as can be seen in, e.g., CD74.

**Figure 3.**
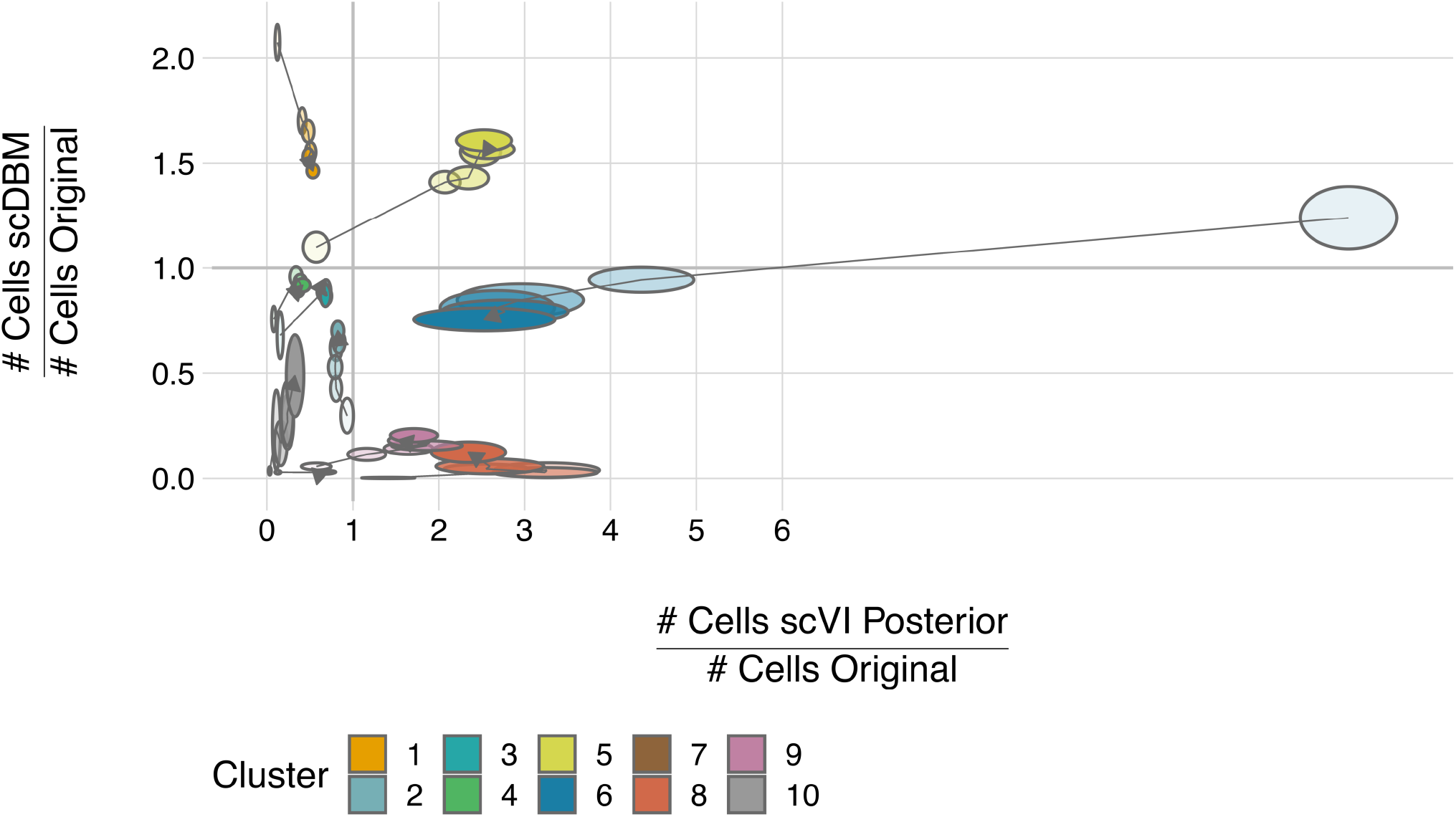
Performance concerning recovering cell-type abundances. Each ellipse represents the number of cells in a specific cluster from one of the generative models divided by the number of cells in the original cluster. scDBM and scVI were trained on a subsample of 384, 768, 1152, 1536, 1920, and 2304 cells (marked by opacity where 384 cells are shown brightest), after which 4182 cells were generated. The y-axis indicates the number of cells per cluster generated by the scDBM divided by the number of cells per cluster in the original data. Hence, if the proportion is higher than one, scDBM overestimated the number of cells in that cluster. The x-axis exhibits the same for samples from *scVI*_*posterior*_. Each bubble is shown six times (for each pilot data set size), and the arrows indicate the change in the ratios. Additionally, the width of ellipses shows the standard deviations of cluster proportions for the 30 subsamples. We have divided the number of cells for each synthetically generated cluster by the number of cells in the corresponding original cluster. If one of the models under study generated the correct number of cells for a given cluster, then this ratio would be 1 and would lie on one of the grey lines.

**Figure 4.**
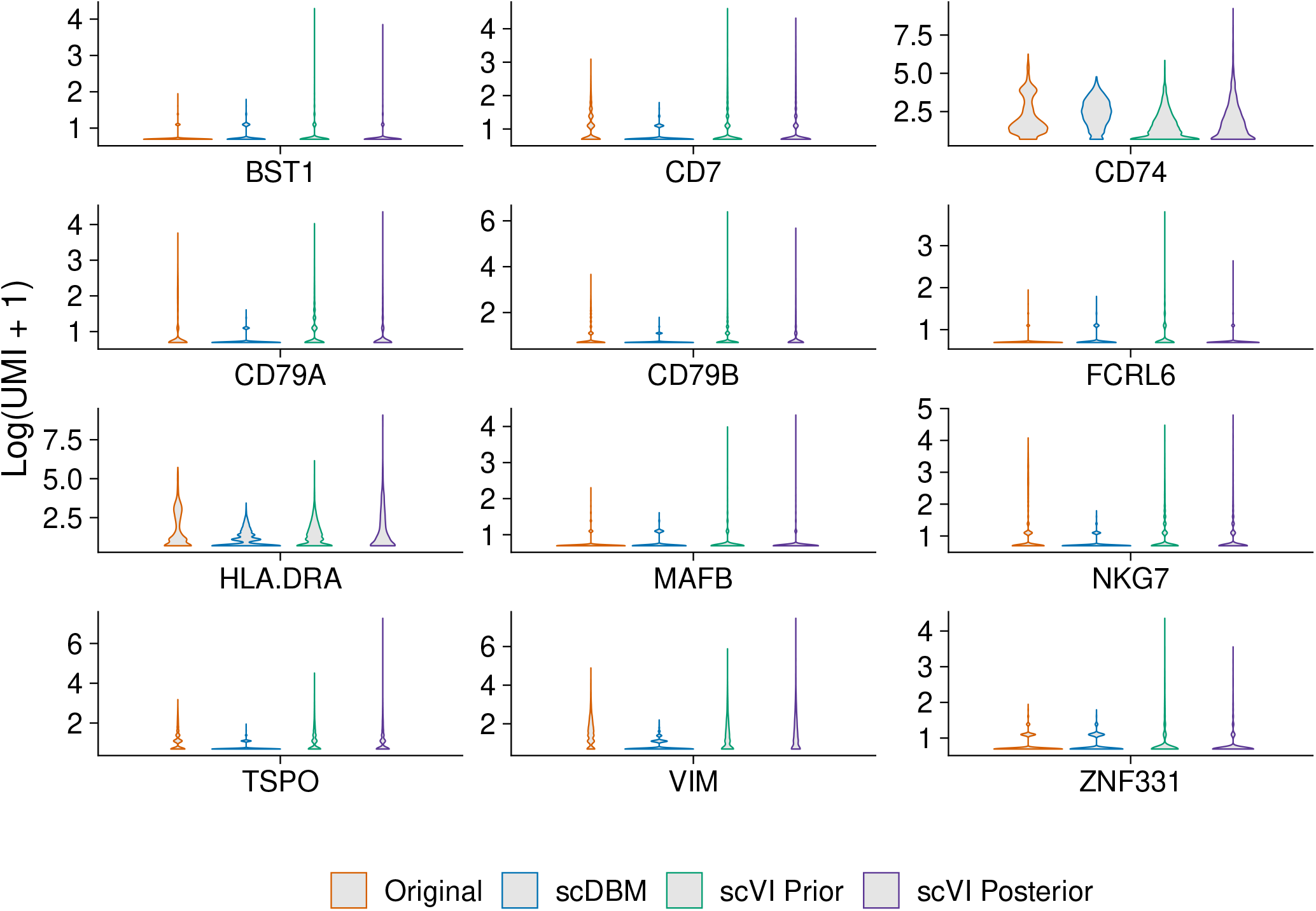
Univariate distributions of expression values for exemplary genes, as generated by scDBM and scVI when trained on 384 cell pilot data subsampled from the PBMC4k data set, compared to the original data.

## Discussion

In this paper, we have investigated the quality of synthetic scRNA-seq data from deep generative models. We looked at situations where we want to draw conclusions from small amounts of data to larger, ground-truth data. This might be relevant, e.g., when planning single-cell RNA-sequencing experiments. To investigate the quality of the generated data, we have relied on three approaches. We used the Davies-Bouldin Index to compare the clusterings’ quality on the synthetic data with the original data. We also looked at how the different models behave in response to varying cluster sizes, and finally, we visually examined the synthetic data using univariate gene distributions.

We subsampled parts of the original dataset to mimic a pilot data scenario. Next, we trained the deep generative models on these subsamples and generated synthetic observations in the original data size. For this, we used scDBM and scVI, where we draw samples from both the prior and posterior distribution for the latter model. In particular, when looking at small datasets, which may be subject to sampling artifacts, it is advisable to draw from the prior distribution, instead of the posterior, in scVI. Furthermore, in such scenarios, Markov chain Monte Carlo methods might have an advantage over variational inference, which is mainly reflected in the lower variability of scDBM. The results show that larger data sets might be necessary to generate synthetic data with proper cluster proportions.

If conclusions are drawn from a small sample to a larger data set, it is important that the pilot sample is representative. In the example of planning an experiment, the pilot study sample may subsequently be included in the main study. Still, care should be taken to ensure that the experimental settings are not changed to ensure representativeness^17^. Studies on the generalization ability of deep generative models already exist^18^, but they have not yet been extended to the application of single-cell transcriptomics data. Doing so is outside the scope of this paper.

Overall, it is a great challenge to infer from a few observations to larger datasets and, depending on the field of application, to monitor the corresponding quality characteristics. Models that specifically target the discovery of rare events could likely provide further performance improvements. Finally, we are confident that deep generative models have great potential for generating synthetic datasets. In particular, these methods could mean an improvement in the planning of future experiments.

## Methods

### Single-Cell Variational Inference

Lopez et al.^7^ proposed a method called single-cell variational inference (scVI), which utilizes the structure of VAEs to encode the transcriptome onto a lower-dimensional representation from which the input is reconstructed. Just as the scDBM, scVI is also based on the (zero-inflated) negative binomial distribution^7^. The model comprises two components, the encoder and the decoder parts of the network. Lopez et al.^7^ use four neural networks for encoding the size factors and the latent variables using the variational distribution *q*(*z*_*n*_, *l*_*n*_ *x*_*n*_, *s*_*n*_) as an approximation to the posterior *p*(*z*_*n*_, *l*_*n*_ *x*_*n*_, *s*_*n*_), where *z*_*n*_ is a low-dimensional vector of Gaussians, *l*_*n*_ is a one-dimensional Gaussian encoding technological differences in capture efficiency and sequencing depth, *x*_*n*_ is the vector of observed expressions of all genes of cell *n*, and *s*_*n*_ describes the batch annotation for each cell^7^. The variational distribution can be written as:

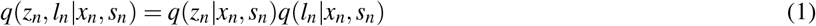

Therefore, the variational lower bound is:

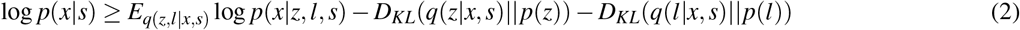

The probabilistic model of scVI is based on a gamma-Poisson mixture. It starts by sampling from the latent space, a standard multivariate normal distribution, which is then fed into a neural network—together with the batch annotation. The neural network then learns the mean proportion of transcripts expressed across all genes. The output is used to sample from a gamma distribution together with the inverse dispersion *θ*_*m*_. The model accounts for technical effects by incorporating a library size scaling factor which, in combination with the gamma-distributed samples, is used to sample from a Poisson distribution. This mixture of the gamma and Poisson distribution is equivalent to the negative binomial distribution^7^. scVI additionally learns a neural network to account for technical dropouts.

Observations are generated from the scVI approach by using original data as input and then sampling from the posterior distribution *p*(*z|x*). A straightforward approach for generating more samples than were used during training is to create (multiple) copies of the original data. For example, for scVI trained on 384 cells, we sampled from the model seven times and stacked the resulting samples together to make inference about a larger number of cells. As an alternative, we adapted scVI to sampling from the prior distribution *p*(*z*) instead of the more common sampling from the posterior *p*(*z|x*). To do that, we changed the inference procedure to sample latent *z* from *Normal*(0, 1) and library sizes from *Normal*(*l*_*μ*_, 1).

### Single-Cell Deep Boltzmann Machine

We adapted deep Boltzmann Machines (DBMs), an unsupervised neural network approach with multiple hidden layers^3^, to the negative binomial distribution. Specifically, we employ the exponential family harmonium framework^19^ that allows restricted Boltzmann machines (RBMs), the single-hidden layer version of DBMs, to deal with any distribution from the exponential family as input. This framework was further extended and simplified by Li et al.^20^.

We use a parametrization of the negative binomial probability mass function that has been suggested by Risso et al.^21^:

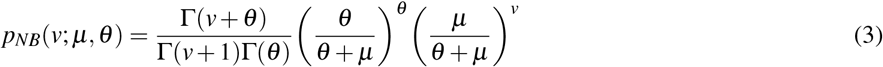

The mean of the distribution is denoted as *μ*, the variance is given by *μ* + *μ*^2^*/θ*, and *θ* is the inverse dispersion. Γ denotes the gamma function.

For simplicity, we describe a three-layer DBM where the visible layer corresponds to an input of unique molecular identifier (UMI) counts for M genes, which can be modeled by a negative binomial distribution^22^. The first and second hidden layers are denoted as *h*^(1)^ and *h*^(2)^, respectively.

Following Li et al.^20^, we define the energy function of the state *{x, h*^(1)^*, h*^(2)^*}* as:

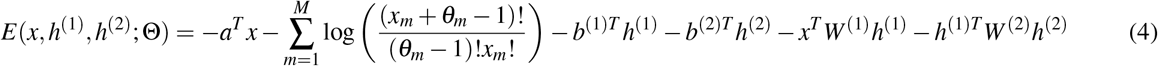

Here, *a*, *b*^(1)^, and *b*^(2)^ are the bias terms of the first, second, and third layer, respectively. Furthermore, *W* ^(1)^ and *W* ^(2)^ denote the weight matrices connecting the layers. Hence, Θ = (*θ, a, b*^(1)^*, b*^(2)^*,W* ^(1)^*,W* ^(2)^) are the model parameters. Therefore, the probability of the visible vector is defined as:

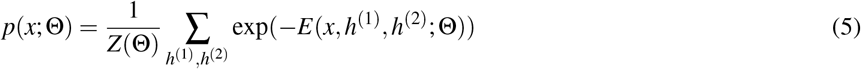

*Z*(Θ) is the partition function which is typically intractable^3^. According to this, the conditional distributions over the visible and the two sets of hidden units are given as:

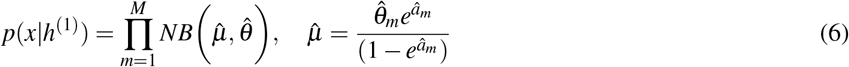

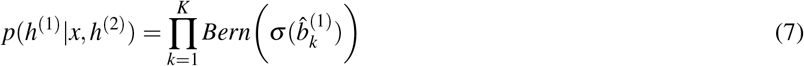

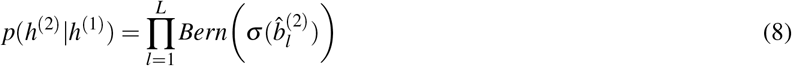

Where 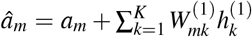 represents the estimate for the visible bias of UMI counts per gene *m* (*m* = 1*,…, M*) and the bias of the first and second hidden layer correspond to 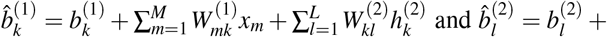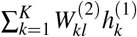, where *k* = 1*,…, K* and *l* = 1*,…, L* indicate the hidden nodes in the first and second hidden layer, respectively. The sigmoid activation function is denoted as *σ*. Training of the scDBMs via stochastic gradient descent can be performed just as for standard DBMs. For a detailed description, see Salakhutdinov and Hinton^3,23^.

After training, synthetic observations can be generated by Gibbs sampling. It can be shown that Gibbs sampling produces asymptotically exact samples, which leads to more accurate results as compared to VAEs^15,24^. This comes at the cost of a higher computational burden, which might be acceptable in small sample scenarios. In contrast, scVI uses variational inference, which scales to scenarios with millions of observations but does not have the advantage of generating exact samples^15^.

#### Estimating the Dispersion Parameter

For the negative binomial distribution, we also need to determine values for the inverse dispersion parameter of each gene which is notoriously difficult^25^.

We use a regularized Fisher scoring algorithm^26^ to estimate the inverse dispersion parameter *θ*_*m*_ for each gene *m*. For this, we use the log-likelihood function of the negative binomial probability mass function (**Equation (3)**) indicated above. The Fisher scoring algorithm can be derived using a two-term Taylor expansion of the score function, the first derivative of the log-likelihood, at the initial choice of the inverse dispersion 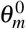^27^. To stabilize estimates of the inverse dispersion parameters, we add 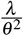 as a regularization term to the log-likelihood, which results in the following scoring algorithm:

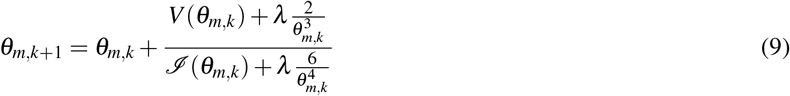

Here, *V* (·) is the score function, ℐ (·) denotes the Fisher information matrix, *λ* is the regularization parameter, and *k* is the current iteration step.

The inverse dispersion parameter *θ*_*m*_ corresponds to the amount of heterogeneity between cells, where a smaller *θ*_*m*_ indicates more heterogeneity. Recall that the negative binomial variance is defined as *μ* + *μ*^2^*/θ*. Due to the regularization term in our model, smaller *θ*_*m*_ are subject to larger regularization. This ensures that we learn the baseline variability between cells, without deflating the estimates of the inverse dispersion due to, e.g., differences between clusters of cells or excess zeros.

#### scDBM Training

By combining the scDBM with Fisher scoring, we can estimate all model parameters Θ = (*θ, a, b*^(1)^*, b*^(2)^*,W* ^(1)^*,W* ^(2)^). In the first step, we initialize all parameters at some reasonable values and learn only a subset of Θ, namely, (*a, b*^(1)^*, b*^(2)^*,W* ^(1)^*,W* ^(2)^). Hence, the inverse dispersion is fixed. After a predefined number of epochs, say five, we use the regularized Fisher scoring algorithm to estimate the inverse dispersion parameter 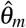 and plug the new estimate into the scDBM. Accordingly, all parameters of the scDBM are refined after a fixed time, e.g., every five epochs.

During training, biases and weights of the network have to be constrained, where *a*_*m*_ = *min* {*a*_*m*_, −*ɛ*} with *ɛ* = 10^−10^ and *w*_*m,k*_ = *min* {*w*_*m,k*_, 0}. This is done because we use the natural form of the exponential family and hence *a*_*m*_ is used in logarithmic scale^20^.

### Evaluation of Synthetic Data Quality

The overall approach taken here for evaluating the quality of generated synthetic observations is illustrated in **Figure 1**. Specifically, a relatively large original dataset is used as ground truth data, and deep generative approaches are tasked with generating synthetic data based on pilot datasets drawn from the original data. We consider Seurat clustering^28,29^ on the UMAP representations^30^ of the original data as a typical data analysis workflow, which provides ground truth cluster labels for the original data. When subsequently assessing synthetic data, each generated observation is assigned the cluster label of the nearest original observation, as determined by Euclidean distance. If a generative approach can provide synthetic observations very close to the original observations, these cluster labels will correspond to a reasonable clustering solution also in the synthetic data. Thus, we can evaluate the quality of the synthetic data by calculating summary statistics for the clusters in the synthetic data, and compare them to cluster statistics from the original data.

Specifically, we use the Davies-Bouldin index (DBI)

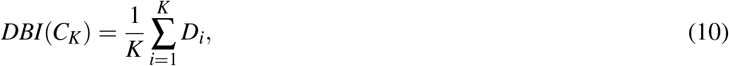

Where

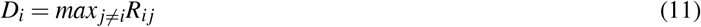

with between-cluster similarity

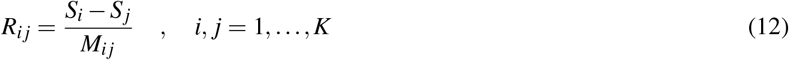

the distance between cluster centroids

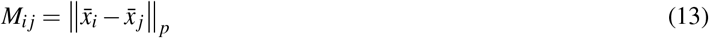

and within-cluster dispersions

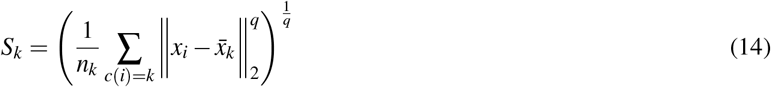

where we set *p* = *q* = 2. Consequently, a small DBI indicates homogeneous and well-separated clusters^31,32^.

To examine whether the models learn to adequately represent frequencies of different cell types, we also compare the number of cells per cluster in the original data and the synthetic observations.

It should be noted that an in-depth evaluation of samples, instead of comparing model fit based on the log-likelihood, is indispensable because it was shown that comparing deep generative models based only on the log-likelihood can be misleading. In particular, even when log-likelihood is low, the quality of generated samples can be good and vice-versa^9^. In contrast, we focus on properties, such as cluster quality, which are important for experimental design.

### Tuning Deep Generative Models

Since we want to investigate the quality of the generated data with the least possible influence of the hyperparameters of the individual models, we keep the architectures of the networks rather simple. The learning rate of the scDBM has to be set relatively small. This is because the reconstruction error can get very large for high expression values. Hence, the corresponding weights will get a very big learning signal^33^. It follows that we also slightly increased the number of epochs. Other than that, we keep the hyperparameter settings largely the same for all models (Supplementary Table 1). We tuned the hyperparameters of the respective models only once per dataset. Hence, we trained the models independently of the size of the dataset. In reality, we would of course tune the networks explicitly for the dataset at hand and most likely achieve better performance. However, tuning each model by hand would be unfeasable in this setting (30 subsamples, 6 dataset sizes, 3 models — 540 trainings in total). We split the data into random train and test subsets with 0.7 being the proportion of the data included in the trainingset.

For scVI, we used the default ReLU activation functions for hidden layers and the sigmoid activation function for hidden layers in scDBMs. We stick to the default parameters for the architecture of scVI. Therefore we use one hidden layer for both the encoder and the decoder network. Furthermore, the dimensionality of the latent space defaults to 10. We also chose two hidden layers for the scDBM and set the dimensionality of the latent space between two to four, depending on the dataset.

### Data Description and Processing

We evaluate the performance of the two scVI variants and the scDBM approach on three typical datasets. First, a 10x Genomics dataset containing peripheral blood mononuclear cells from a healthy donor is considered^34^. We preprocessed the data following Amezquita et al.^35^, after which 4182 cells and 1000 highly variable genes were left for downstream analysis. We refer to this dataset as *PBMC*4*k* throughout this work.

Second, analyses are performed on a data set of neuronal subtypes in the mouse cortex and hippocampus, where Zeisel et al.^36^ sequenced 3005 cells from male and female juvenile mice. We specifically consider data from 2816 cells and 1816 highly variable genes which were left after preprocessing^35^. We refer to this dataset as *Zeisel* throughout this work.

Additionally, we demonstrate the performance on a currently unpublished scRNA-seq dataset from the hippocampus of three embryonic (E16.5) mice processed with the CEL-Seq2 protocol^37,38^. The unnormalized count matrix contained 3808 cells, and we selected the 1500 most highly variable genes for downstream analysis. We used scran and scater^39,40^ for pre-processing. We refer to this dataset as *Hippocampus*4*k* throughout this work. The results for *Zeisel* and *Hippocampus*4*k* can be found in the supplementary material.

### Implementation

The scDBM implementation is based on the Julia package ‘BoltzmannMachines.jl’^41^ and extends the packages’ scope to scRNA-seq data which is available at https://github.com/MTreppner/scDBM.jl.

Furthermore, we used the Python implementation of scVI (https://github.com/YosefLab/scVI), which we adapted to be able to sample from the prior distribution.

## Acknowledgements

The work of M.T. and A.S.B has been funded by the Deutsche Forschungsgemeinschaft (DFG, German Research Foundation) - 322977937/GRK2344. The work of M.H. has been supported by the Federal Ministry of Education and Research in Germany (BMBF) project ‘Generatives Deep-Learning zur explorativen Analyse von multimodalen Omics-Daten bei begrenzter Fallzahl’ (GEMOLS: Generative deep-learning for exploratory analysis of multi-modal omics data with limited sample size, Fkz. 031L0250A). The work of SL has been supported by the BMBF in the MIRACUM project (Fkz. 01ZZ1801B).

## Author contributions statement

H.B. conceived the methods, M.T. conducted the analyses, implementations and wrote the manuscript, M.H. and S.L. contributed in analysis and results interpretation. A.S.B and T.V. performed wet-lab experiments for the Hippocampus dataset and proofread the MS. All authors read and approved the final manuscript.

